# A Composite Endomembrane Suspension Governs Cytoplasm Rheology

**DOI:** 10.64898/2026.07.20.739612

**Authors:** M.I. Arjona, A. Khosravanizadeh, C. Municio-Díaz, M. Mioche, S. Dmitrieff, J. Sallé, A. Marteil, C. Pioche-Durieu, L-L. Pontani, E. Wandersman, N. Minc

**Author notes:** Correspondence to : M.I.A; L.L.P; E.W and N.M.

## Abstract

The cytoplasm of eukaryotic cells is populated by dense disordered suspensions of filamentous and granular endomembranes, yet how they contribute to the mechanical behavior of the cell interior remains unknown. We combined active micro-rheology, cell-like encapsulation and simulations to study the material properties of marine egg extract fractions enriched in distinct endomembrane components. We characterized the cytoplasm as a composite suspension made of yolk granules interspaced by sheets and tubules of endoplasmic reticulum (ER) bathed in cytosolic fluid, that occupies ∼38% of cell volume. Remarkably, while isolated cytosol, yolk or ER fractions had characteristics of Newtonian fluids, their combination yielded the emergence of viscoelasticity and glass-like dynamics closely resembling that of *in vivo* cytoplasm. Our data suggest that the ER acts as a sterically excluding backbone that drives the formation of load-bearing yolk flocculation structures to endow the cytoplasm with solid-like properties at volume fractions far below random close packing. This work establishes a generic framework to understand the material properties of composite endomembrane suspensions, and delineates a novel strategy by which eukaryotic cells may tune the physical state of their cytoplasm.

---

The cytoplasm is an active water-based medium, crowded with macromolecules like proteins and mRNAs, as well as larger endomembrane compartments and cytoskeletal networks. These components may behave as dense heterogenous coarse suspensions or as percolated polymer gels, to endow the cytoplasm with complex rheological properties, that have fundamental implications for cellular and multicellular biology^1–8^. Accordingly, recent physical descriptions of the cytoplasm have invoked frameworks of soft colloidal glasses approaching the jamming transition^6,9,10^, dense porous media^11,12^, or polymeric networks^13–15^. The density of components occupying cytoplasm volume is one fundamental parameter that regulates its material properties. This is best exemplified by alterations in cytoplasm viscoelastic constants in response to osmotic compression or relaxation that directly modulate intracellular densities^9,11,16^. For instance, at the macromolecular ∼10-100 nm scale, proteins, RNA and complexes like ribosomes may occupy volume fractions of ∼10-15%^7,8^. Although far below random close packing of hard spheres, such colloidal crowding is now well recognized to generate enhanced microscopic viscosity, restricted diffusion and local caging effects^4,6,7^.

At macroscopic scales above the micron, relevant to cellular spatial organization and morphogenesis, the mechanics of the cytoplasm has been most often linked to the presence of cytoskeleton polymer networks^1,5,13,15,17,18^. However, in many cell types, depolymerization of the cytoskeleton may only have limited impact on cytoplasm viscoelastic behavior^13,15,16,19,20^. This suggests that cytoplasm material properties could be governed by other elements than cytoskeleton networks, and rather tuned by the cytoskeleton, during cellular transitions such as the cell cycle or differentiation^13,15,21,22^. At this scale, the eukaryotic cytoplasm is packed with endomembrane compartments, in the form of lipid droplets, vesicles, tubules or sheets. Depending on cell types, these may occupy volume fractions of up to 20-50%, with large organelles like the endoplasmic reticulum (ER) taking as much as 7-10% of intracellular volumes on its own^23,24^. To date, however, how the packing of endomembrane components influences cytoplasm physical properties has been left unexplored, given the difficulty of removing or manipulating these compartments in living cells without compromising cell survival.

Here, using sea urchin eggs, as a powerful model for studies of cytoplasm rheology ^2,16,19,21^, we employed a bottom-up approach to isolate endomembrane suspensions, and measure their individual contribution to cytoplasm viscoelasticity. Our findings support a model in which the synergistic combination of a colloidal suspension of yolk lipid droplets intertwined by ER filaments, yields the formation of a composite material with viscoelastic properties and glassy dynamics, that recapitulate *in vivo* cytoplasm physical behavior.

## A composite endomembrane suspension fills the cytoplasm of living cells

Sea urchin unfertilized eggs are arrested in interphase of the cell cycle, and lack organized cytoskeleton networks, featuring only sparse F-actin filaments and nearly no microtubules in their cytoplasm. Yet, their cytoplasm was previously shown to behave as a Jeffreys viscoelastic fluid, with a macroscopic shear viscosity of ∼1 Pa.s and bulk elastic modulus of ∼0.5 Pa, of similar magnitude as in other cell types^3,13,16,20^. Importantly, F-actin disruption only marginally affected viscoelastic constants^16^, suggesting that a cytoskeletal polymer-based physical description of the cytoplasm may not be relevant^5,13,15,17^. In order to characterize macroscopic components that crowd the cytoplasm as potential mechanical regulators, we used Serial Block Face Scanning Electron Microscopy (SBF-SEM) to image the whole volume of these ∼100 µm sized cells, with an axial resolution of ∼50 nm^19^. This revealed that the cytoplasm is homogeneously packed with endomembranes, comprising yolk granules, mitochondria and acidic organelles (lysosomes and endosomes), as well as tubules and sheets of the ER. Using supervised-learning to automatically segment and classify different components, we quantified that yolk granules, forming a monodisperse suspension, occupy a volume fraction of ∼26 % being the most abundant crowders at this scale, while mitochondria occupy only ∼2-3%, and other vesicles less than 1%. Interestingly, this analysis revealed that the ER network, in the form of tubules and sheets, filled interspaces between vesicles and granules taking on its own a volume fraction of ∼7-8% (Fig. 1a and 1d and Movie S1). Therefore, this cytoplasm is filled with a composite endomembrane suspension made of colloidal droplets intertwined by filamentous-like ER that occupy altogether ∼38% of the cell volume.

**Figure 1.**
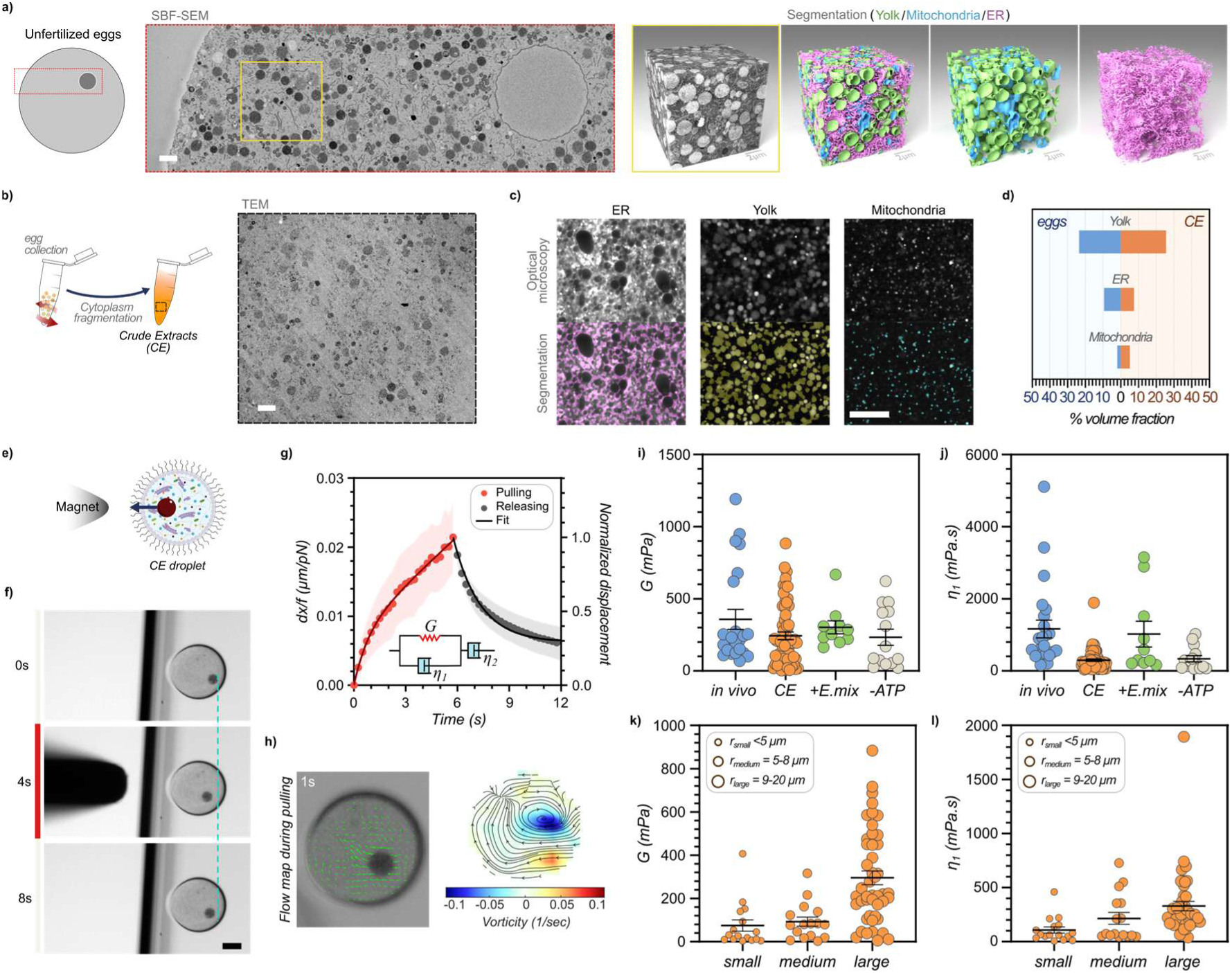
*In vitro* reconstitution of sea urchin eggs cytoplasm crowding and rheology. **a**, Unfertilized Sea urchin egg section imaged with SBF-SEM and pixel classification of different organelles in a 3D reconstruction of a cytoplasm cube portion. Scale bars: 2 µm. **b,** Schematic of crude extracts (CE) generation from sea urchin egg fragmentation and transmission electron microscopy image of CE. Scale bar: 2 µm. **c,** Spinning disk confocal images of CE stained for ER (DilC18), yolk (Nile Red), and mitochondria (Mitotracker), and image segmentation used to quantify volume fractions. Scale bar: 10 µm. **d,** Quantification of the yolk, ER, and mitochondria volume fractions in eggs and in CE. **e,** Scheme of the crude extracts encapsulated in droplets together with magnetic beads displaced with magnetic tweezers. **f,** Time-lapse images of magnetic bead displaced inside a CE filled cell-like compartment by an external magnetic force and recoiling when force is removed. Scale bar: 50 µm. **g,** Displacement curve scaled by applied force during the pulling phase, and normalized relaxation curve after force removal for CE encapsulated in droplets. Fits correspond to that of a Jeffreys model which consists of a spring, *G*, and a dashpot, *η_1_*, in parallel and a second dashpot, *η _2_*, in series (n=20). **h,** (left) Fluid flow vector field of CE during force exertion, and (right) streamlines averaged on a duration of 2s overlaid on a vorticity map obtained by PIV. **i,j,** Elastic (i) and viscous (j) moduli and corresponding viscoelastic timescale (k) obtained from fitting creep curves with Jeffreys’ model for in vivo cytoplasm^16^(n=23), CE (n=62), CE supplemented with an energy mix solution (n=10), and ATP-depleted CE solution (n=14). **k,l** Crude extract viscoelastic parameters as a function of object sizes (n=16, n=16, n=42, for small, medium and large objects, respectively). Error bars correspond to +/-s.e.m, and the shades in 1h correspond to +/- s.d.

## *In vitro* reconstitution of cytoplasm crowding and material properties

To understand if and how endomembranes may promote cytoplasm viscoelastic behavior, we set out to reconstitute the cytoplasm using a bottom-up approach^25^. We first generated complete undiluted crude extracts, and confirmed by both electron and fluorescent microscopy that these contained endomembranes at similar volume fractions as the *in vivo* cytoplasm (Fig. 1b-d, Extended Fig. 1a-b and Movie S2). To reconstitute cellular boundaries and perform active rheological measurements, we encapsulated extracts together with magnetic particles, in ∼100-200µm cell-like compartments suspended in oil and stabilized by surfactants (Fig. 1e and Extended Fig. 1c-e)^26,27^. Remarkably, when submitted to calibrated magnetic-based forces, generated by approaching a magnetized tip, extracts exhibited a viscoelastic response typical of a Jeffreys material, similar to that observed *in vivo*^2,16,19^. This response encompasses a short-term viscoelastic behavior characterized by a viscous dashpot, *η*_1_ in parallel with an elastic spring, *G*, and a long-term fluidization response characterized by a second viscosity, *η*_2_. This behavior was evident from the partial recoil of magnetic beads when the force was released, that reflects partial dissipation of stored elastic energy, as well as viscoelastic flows, recirculations, and endomembrane deformations induced by the moving particle, that indicates that cytoplasm extracts deform elastically at short time-scales but flow at longer time-scales (Fig. 1e-h and Movie S3-S4).

Fits of the creep response, allowed to compute a bulk elastic modulus G= 243.3 +/- 26.7 mPa (+/- indicate s.e.m) and a short-term shear viscosity η_1_ = 299.3 +/- 35.1 mPa.s, yielding a viscoelastic time-scale of 3.99 +/- 1.08 sec (Fig. 1i,j and Extended Fig. 2a). Importantly, these mechanical constants were of similar magnitudes as that previously measured for the *in vivo* cytoplasm^16^. However, we noted a marked reduction in the values of viscous moduli in extracts, indicating a potential loss of endomembrane turn-over as compared to real cells. Accordingly, supplementing extracts with an energy mix containing both ATP and GTP, to promote ER membrane fusion and turnover^28,29^, increased viscoelastic values to G= 302.2 +/- 45.2 mPa and η_1_ = 1023.1 +/- 358.7 mPa.s, closer to that found *in vivo* (G= 356.9 +/- 68.9mPa and η_1_ = 1163.0 +/- 244.6 mPa.s). In contrast, ATP depletion caused ER membranes to partially collapse, concomitant with a reduction in extract viscoelastic constants^9^ (Fig. 1i, and Extended Fig. 2b,c). F-actin filaments were also present in extracts with similar appearance but a reduced density as compared to real cells, and their disassembly by cytochalasin D only caused a small reduction in viscoelastic constants, consistent with measurements *in vivo*^16^ (Extended Fig. 2e-h). These filaments also depolymerized when ATP was depleted, potentially contributing to the reduction of viscoelastic constants in this condition (Extended Fig. 2c). Finally, by probing magnetic particles of different sizes, we recapitulated size-dependent viscous and elastic behavior previously reported *in vivo*, indicating that most representative pore sizes of the cytoplasm may be kept intact in extracts (Fig. 1k,l)^16^. We conclude that crude extracts provide a *bona fide* representation of *in vivo* cytoplasm macroscopic crowding and material properties.

## The cytoplasm exhibits physical characteristics reminiscent of colloidal gels

Colloidal crowding models of cytoplasm mechanics have been supported by varying volume fractions and probing impact on viscoelastic responses^9^. In cells, this is commonly achieved through osmotic perturbations that alter water content, cell volume and thus volume fractions^9–11^. We explored these aspects by serially diluting or over-concentrating crude extracts (Fig. 2a-c). At dilutions below 30%, extracts behaved as Newtonian viscous fluids with nearly no detectable elastic response, and shear viscosities typically ∼5X smaller than undiluted extracts. However, at concentrations above ∼40-50%, we detected the clear emergence of elastic behavior, accompanied by a marked increase in shear viscous moduli. Both elastic and viscous moduli kept increasing above this critical concentration in dose-dependence with extract concentration (Fig. 2a-c; Extended Fig. 3a-c). This indicates that the cytoplasm may transit from liquid-like to solid-like at an estimated critical volume fraction in all endomembranes of ∼15-20%, and only ∼10% in yolk droplets, far below jamming which is achieved at a random close packing density of ∼64% for monodisperse spheres^30^. These findings indicate that a simple colloidal description, in which elastic behavior emerges at random close packing may not simply explain cytoplasm mechanical behavior^30^. Accordingly, by fitting with a power-law the evolution of the elastic modulus as a function of yolk particle fraction in the serially diluted extracts above a critical volume fraction φ_c,_ as G=(φ/φ_c_–1)^ν^, we computed critical exponents ν, ranging between ∼2 and 3 (depending on the exact chosen critical volume fraction). These exponents are much larger than the ½ exponent expected for colloidal jamming above random close packing^30^, and rather closer to those derived for colloidal gels^31^ (Fig. 2d). In such gels, weak attractive forces among particles lead to the formation of percolated particle flocs that serve as load-bearing structures to promote solid-like behavior at low volume fractions^32^. Imaging of yolk particles in crude extracts or in EM images of real cells confirmed a largely inhomogeneous distribution of yolk particles, and multiple clear instances of connected flocculated structures (Fig. 1a and 2e,f). Together, these analyses suggest that composite membrane networks may exhibit some characteristics reminiscent of colloidal gels to endow solid-like behavior to the cytoplasm at low volume fractions.

**Figure 2.**
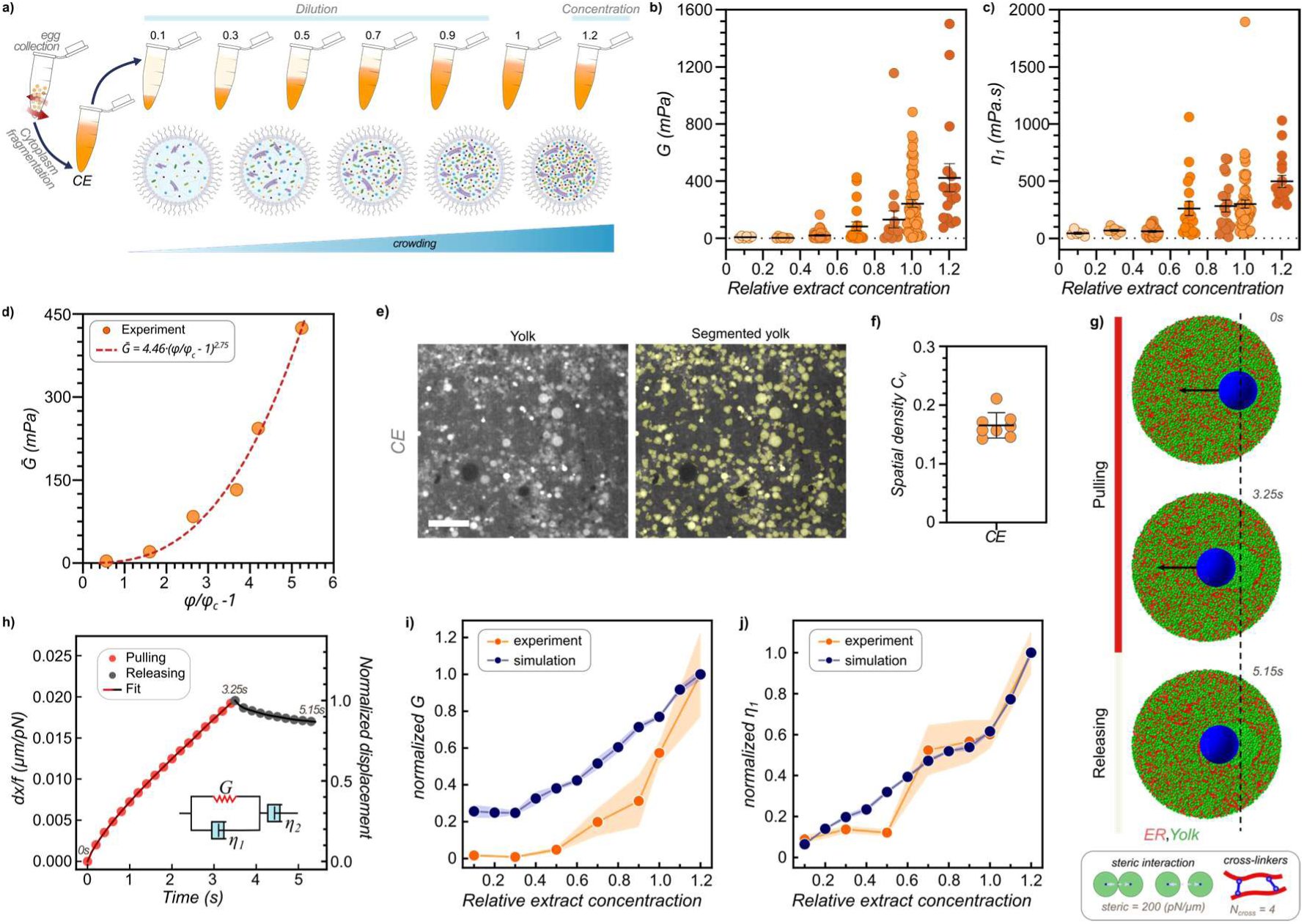
Impact of extract crowding on viscoelastic behavior. **a**, Pure CE solution obtained from unfertilized Sea urchin eggs was brought to different concentration by diluting or concentrating initial undilute CE, and encapsulated in cell-like compartments to monitor viscoelastic behavior. **b,c,** Elastic and viscous moduli as a function of relative extract concentration (1 corresponds to undiluted CE) (n=6, n=7, n=30, n=19, n=19, n=62, and n=17, respectively). **d,** Mean values of the elastic modulus plotted as a function of yolk volume fraction above a critical fraction Φ_c_ =0.05 and fitted with a power-law. **e,** Spinning disk confocal images of CE stained for yolk (Nile Blue), and segmented image used to extract particle positions for fractal analysis. Scale bar: 10 µm. **f,** Spatial density coefficient of variation, C_v_, for yolk distribution in CE (n=8); C_v_ vanishes to 0 for a perfectly homogeneous distribution. **g,** Agent-based simulations for creep and relaxation with a probe inside a sphere filled with spherical particles that repulse each other’s (Yolk) and filaments connected by cross-linker (ER) at similar densities as that measured in CE or *in vivo*. **h,** Representative displacement (red) and normalized recoiling (grey) curves for the simulation presented in 2d. **i,j,** Normalized elastic and viscous moduli of the different extract concentration computed from simulation (blue) and experiments (red). For the simulation, dots represent the average of 5 independents simulations, for the experiments they show the average from b) and c). Error bars and shades correspond to s.e.m.

## Agent-based simulations of composite endomembranes recapitulate the emergence of cytoplasm viscoelasticity

To test if this endomembrane composite suspension can generate macroscopic viscoelasticity, we next implemented agent-based simulations. We modeled yolk granules as rigid particles of 1 µm in diameter, and included a steric repulsion term to prevent overlaps. The ER was coarse-grained and modeled as a set of elastic filaments of 0.2 µm in diameter and 6 µm in length, with a bending modulus taken from previous measurements *in vivo*^33^. ER filaments were connected by crosslinkers to percolate the network, and favor weak attractions among filaments. Finally, to simulate creep and relaxation assays as in experiments, we added a large probe (8 µm in radius) actuated by an external constant force. In simulations, the presence of long ER filament bundles tended to exclude yolk particles resulting in flocculation zones as seen in experiments. Furthermore, at volume fractions taken from experimental measurements, these simulations recapitulated a Jeffreys-like viscoelastic behavior, with large-scale deformations of endomembranes as observed in extracts (Fig. 2g-h and Movie S4-S5). Simulated viscoelastic constants also scaled with volume fractions as in serial dilution experiments, although we noted some residual elastic behavior even at low volume fractions, presumably reflecting the more brittle or dynamic nature of the ER in extracts as compared to simulations. Viscoelastic behavior was robust to parameter values for steric repulsion, number of crosslinkers per ER filament and did not grossly change when adding specific connections between ER filaments and yolk granules (Fig. 2i,j and Extended Fig. 4a-c). These results indicate that the mere packing of flexible filaments interspacing a colloidal-like yolk suspension may be sufficient to impart viscoelasticity to the cytoplasm even at low volume fractions.

## Decomposing the mechanical contribution of endomembrane networks

To understand how individual cytoplasmic constituents contribute to its mechanical behavior, we next generated different endomembrane extract fractions (see Material and Methods). We first purified a membrane-free cytosol, that can support physiological instances such as sperm chromatin de-condensation and histone phosphorylation^34^, and characterized its rheology. Cytosolic extracts had characteristics of pure Newtonian fluids with a shear viscosity of 24.05 +/- 2.38 mPa.s, similar to values reported for vertebrate cytosol^25^ (Fig. 3a-d; Extended Fig. 5a-d). Using Einstein’s formula for the viscosity of dilute suspensions^35^ allowed to estimate from this measurement, a volume fraction of suspended macromolecules of ∼10%, in agreement with previous estimates based on typical macromolecule concentration and size in cells ^8^. Therefore, background cytosol which presumably retains a large fraction of native soluble macromolecules, only accounts for ∼8% of the viscosity of complete crude extracts at this macroscopic scale.

**Figure 3.**
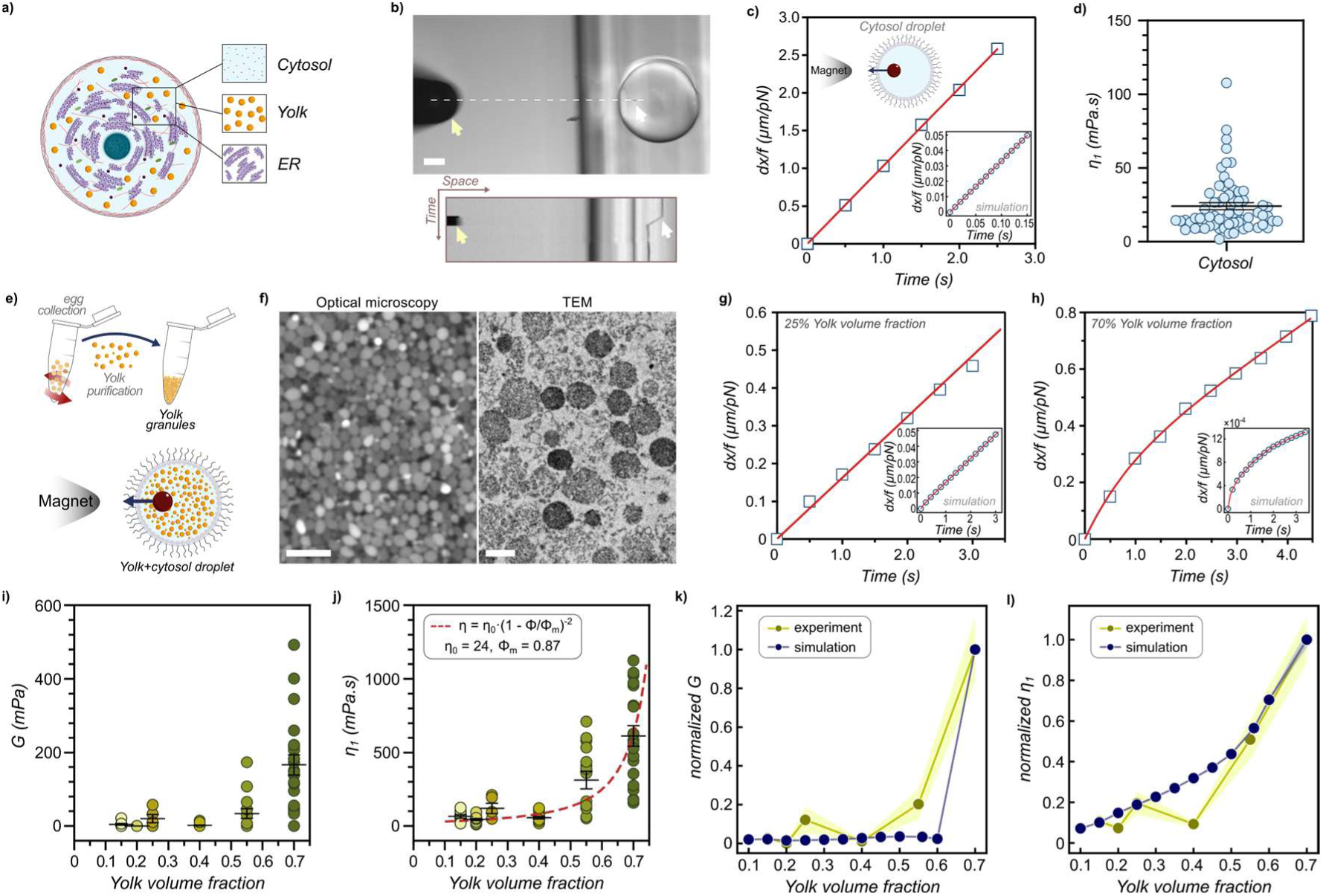
Yolk granules behave as a soft colloidal suspension. **a**, Schematic of the cytoplasm representing different fractions that may contribute to cytoplasm viscoelastic behavior. **b,** Time-lapse image and kymograph of a magnetic bead moved by force in membrane-free cytosol encapsulated in a cell-like compartment. Scale bar: 50 µm. **c,** Representative creep curve *for* membrane-free cytosol encapsulated in a cell-like compartment. Inset correspond to the simulation of a membrane-free fluid of fixed viscosity. **d,** Viscous modulus of the membrane-free cytosol solution (n=64). **e,** Scheme of yolk suspension purification and encapsulation in cell-like compartments for active rheological measurements. **f,** Fluorescent and transmission electron microscopy images of the purified yolk granule suspension. Scale bars: 5 µm and 1 µm, respectively. **g,h,** Creep curve for a diluted Yolk suspension (25%, left), at similar volume fraction as in real cytoplasm, and for a more crowded one (70%, right). Fits correspond to that of a Jeffreys’ model. Insets correspond to simulation results at similar crowding. **i,j,** Elastic and viscous moduli of yolk suspensions in cytosol plotted as a function of yolk granules volume fraction (n=9, n=11, n=5, n=14, n=15, and n=21, respectively). The average values of the viscous moduli are fitted with a Maron & Pierce power law for granular media (j). **k,l,** Normalized elastic (k) and viscous (l) moduli of yolk suspension at varying volume fractions, computed from simulations (blue) and experiments (red). For the simulation, dots represent the average of 5 independents simulations, for the experiments they show the average from i) and j). Error bars correspond to +/- s.e.m.

Next, we isolated yolk granules, and resuspended them in cytosol at varying volume fractions (Fig. 3e). In both experiments and simulations, this suspension exhibited a rheological signature close to that of soft colloidal dispersions (Fig. 3f-l). Indeed, suspension viscosity followed a volume fraction-dependent Maron & Pierce power law with an estimated maximum packing of ∼85% consistent with the deformability of yolk droplets^36,37^ (Fig. 3j). In addition, we noted the emergence of solid-like behavior only above volume fractions of ∼60-70% in agreement with the random close packing of monodisperse spheres (Fig. 3h-j). Importantly, however, at fractions of ∼25% similar to that measured in crude extracts and *in vivo*, this suspension exhibited almost no elastic behavior, and a viscosity of only 39% of complete extracts (Fig. 3g,i,j; Extended Fig. 5e,f and Movie S6). Therefore, these results demonstrate that yolk granules may behave as repulsive deformable colloids, that cannot on their own impart viscoelasticity to the cytoplasm.

We next purified extract fractions enriched in ER alone or in ER and yolk, and resuspended them in cytosol (Fig. 4a-c and Extended Fig. 6a-f). Remarkably, while a pure ER suspension at 7% volume fraction exhibited only weak elastic behavior, and a yolk suspension at 15% exhibited a pure Newtonian signature, a composite suspension combining 7% ER and 15% yolk now had a marked elastic response and a viscosity higher than the sum of viscosities of individual fractions (Fig. 4d-g). Further increasing the density of this reconstituted ER-yolk composite to a level closer to that found in crude extracts or *in vivo* (11% ER-30% yolk), enhanced viscoelastic constants to values near similar to that measured for crude extracts or *in vivo* (Fig. 4h-j and Extended Fig. 7a-d). These results directly evidence a synergistic response created by mixing filamentous and granular endomembranes that may promote solid-like characteristics of the cytoplasm.

**Figure 4.**
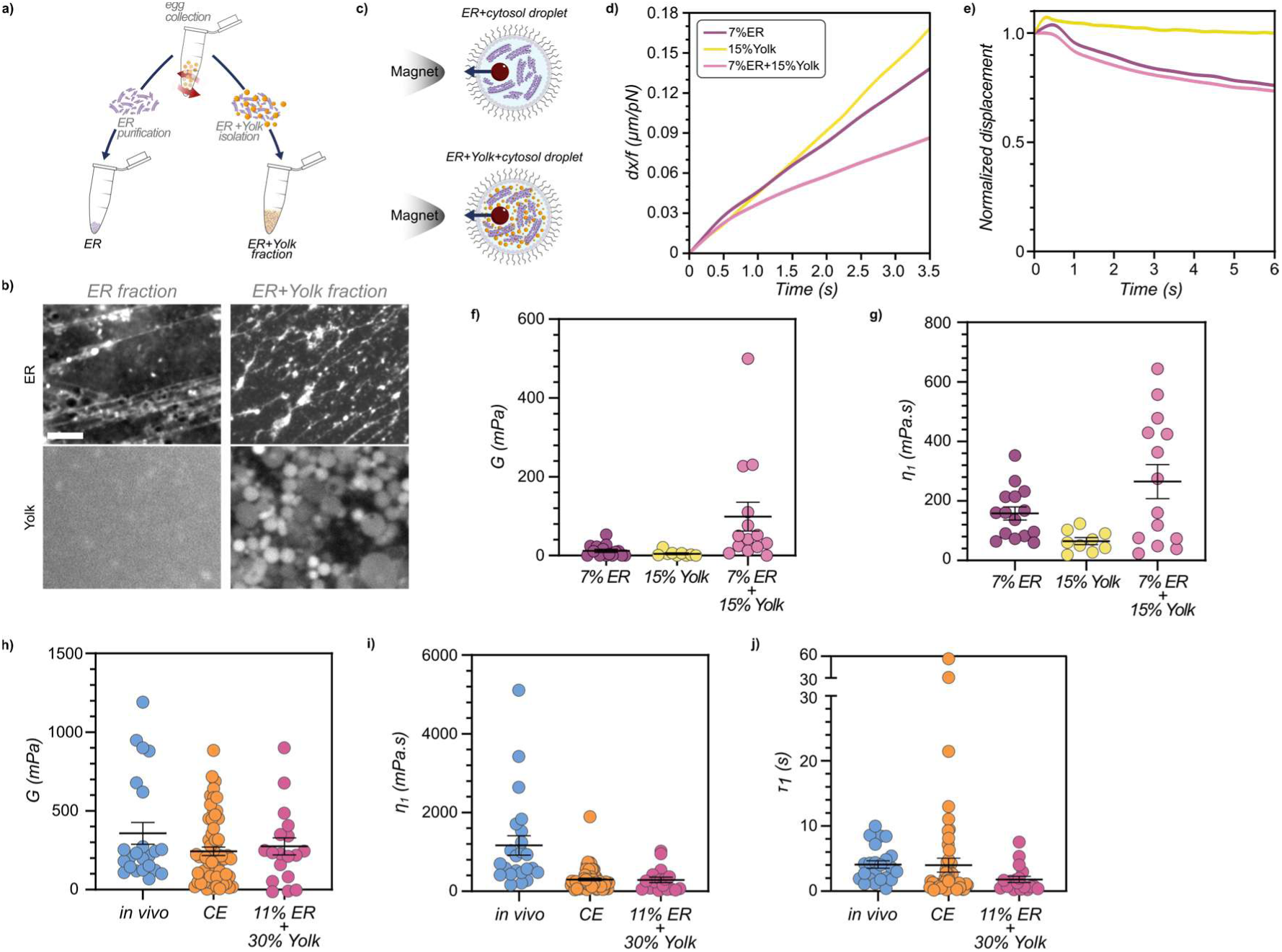
A composite ER-yolk-cytosol fraction recapitulates cytoplasm rheological behavior. **a**, Scheme representing the obtention of pure ER or ER+Yolk fractions. **b,** Confocal images of ER and yolk in pure ER (top) and ER+yolk (bottom) fractions. Scale bar: 10 µm. **c,** Scheme of ER (top) and ER+Yolk (bottom) fractions encapsulation in cell-like compartments for active rheology measurements. **d,e,** (d) Creep (e) and relaxation (e) curves for different endomembrane fractions: 7%ER, 15%Yolk, 7%ER+15%yolk, and CE. **f,g,** Measured elastic (f) and viscous (g) moduli for different endomembrane fractions: 7%ER, 15%Yolk, 7%ER+15%yolk (n=15, n=9, and n=14, respectively). **h,i,j,** Measured elastic (f) and viscous (g) moduli and corresponding viscoelastic time-scale (h) for *in vivo* cytoplasm^16^ CE, and a composite suspension containing 11% ER and 30% Yolk suspended in cytosol (n=23, n=62, and n=18, respectively).

These changes in material properties were also reflected in the diffusive dynamics and in the spatial organization of yolk particles. To track diffusive dynamics, we imaged yolk particles at high frame rates (3.3Hz), and computed their Mean Squared Displacement (MSD) as a function of lag time. This allowed to derive a diffusivity coefficient by fitting the linear part of the MSD curve at short time-scale, as well as estimate a non-linear power exponent α by fitting the curve in logarithmic scale over the whole range of time-scales (Extended Fig. 8). When suspended in cytosol, at a volume fraction of 30%, yolk granules exhibited slight sub-diffusive behavior, with a diffusivity coefficient of 0.0693 +/- 0.0660 µm^2^/s and α=0.77 +/- 0.3, likely reflecting the semi-dilute regime of the suspension. However, when tracked in a composite ER-yolk (11%/30%) suspension or in undiluted crude extracts, they exhibited markedly reduced diffusivity coefficients of 0.0005 +/-0.0010 µm^2^/s and 0.0025 +/- 0.0060 µm^2^/s, as well as lower sub-diffusive α exponents of 0.56 +/- 0.2 and 0.46 +/- 0.2 respectively (Fig. 5a-c). These results reflect pronounced caging effects and restricted mobility caused by the presence of ER filaments interspacing yolk granules. In addition, yolk particles suspended in cytosol were relatively homogeneously distributed reflecting their freely diffusing behavior. In sharp contrast, the presence of an ER network yielded the formation of large connected yolk flocs similar to that found for crude extracts (Fig. 5d-f). These findings indicate that the ER network may serve as a sterically excluding backbone that promotes the formation of flocculated yolk particles structures, driving dynamical arrest and caging effects, marking a transition into a viscoelastic composite reminiscent of a colloidal gel (Fig 5g).

**Figure 5.**
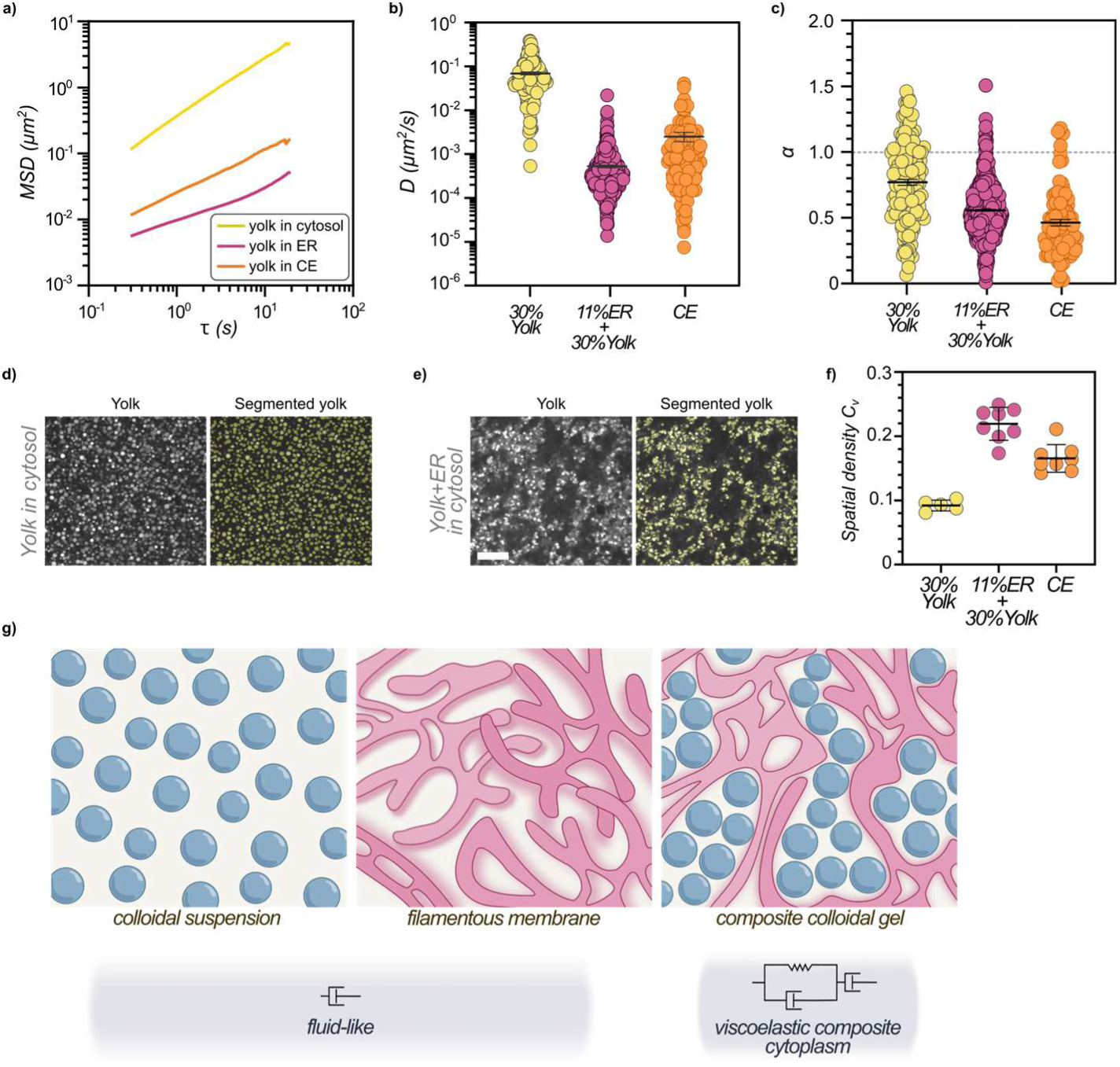
Dynamical caging and restricted mobility in endomembrane composites. **a**, Mean squared displacement (MSD) of yolk granules as a function of the lag-time, *τ*, for a composite suspension of 11%ER+30%yolk in cytosol (purple), a 30% yolk suspension in cytosol (yellow), and CE (orange). **b,c,** Diffusion coefficient, D (b) and scaling exponent *α* (c) obtained from fitting MSD as a function of time-lag as: *MSD* = 4*Dτ*^a^ (n=158, n=763, n=96 corresponding to yolk granules in a 30% yolk suspension in cytosol, in a composite suspension of 11%ER+30%yolk in cytosol, and in CE respectively.) **d,e,** Spinning disk confocal images stained for yolk (Nile Blue), and corresponding image segmentation used to obtain particle position and compute the fractal dimension of a 30% yolk suspension in cytosol (d) and a composite suspension mixing 11%ER+30%yolk in cytosol (e). Scale bar: 10 µm. **f,** Spatial density coefficient of variation, C_v_, for yolk distribution in a 30% yolk suspension in cytosol (n=5), a composite suspension of 11%ER+30%yolk in cytosol (n=8), and in CE (n=8). **g,** Proposed model for how a composite endomembrane suspension can endow the cytoplasm with macroscopic viscoelastic properties: ER filaments may generate depletion-attraction forces among yolk particles, leading to the formation of a composite akin to a colloidal gel, that promote dynamical arrest and the emergence of elastic behavior in the cytoplasm.

## Conclusion

Granular media and colloidal suspensions transit from fluid-like to solid-like, through a dynamical arrest as they reach volume fractions close to random close packing. However, this transition can occur at markedly smaller fractions when particles repulse or attract each other through electrostatic, hydrophobic or hydrodynamic interactions, for instance ^37,38^. As a consequence, composite solutions mixing fluid suspensions with strict Newtonian behavior, like emulsions stabilized by microgel thickener or surfactant micelles linked by co-polymers, can exhibit rich rheological behaviors, with the emergence of elastic, brittle-to-ductile, shear-thinning or shear-thickening behavior^39,40^. Here, we propose, that the macroscopic rheology of bulk cytoplasm is dominated by such a composite system, formed by a suspension of yolk droplets interspaced by ER filamentous membranes. This suspension endows solid-like behavior and glassy dynamics to the cytoplasm at short time-scales below few seconds, but can still flow at longer time scales to promote cellular reorganization.

Our analyses suggest that this suspension may have a set of physical characteristics reminiscent of colloidal gels, with the formation of yolk particle flocs that may serve as connected load-bearing structures to drive an elastic percolation at volume fractions far below random close packing^41^ (Fig. 5g). We show that formation of such structures can be recapitulated by simply mixing ER with yolk particles. Our interpretation of these results is that the ER provides a structural backbone that shapes the granules into a particulate network percolating the cell or cell-like compartment. Similarly to colloidal gels, such system-spanning structures can impart elasticity to the cytoplasm even at low packing fractions, much like in colloid-polymer mixtures^42,43^. However, whether specific dynamic interactions among ER membranes or particular topological properties of its network underlie or modulate such transition remain fundamental questions for the future. Recent findings in single celled eukaryotes suggest, for instance, that ER topological entanglements around large vacuoles may dampen cytoplasm deformations^44^. We also observe some instances of entanglements in EM images (Fig. 1a), suggesting these could potentially also contribute to solidify the composite.

Several evidences suggest that crude extracts retain some ATP-fueled activity inherited from cells. These include the general active dynamical appearance of endomembranes in extracts (Movie S2), the presence of F-actin filaments which require ATP for their assembly (Extended Fig. 2a-b and 2d), and the diffusivity of yolk granules that we estimate to be ∼2-3X higher than that expected from pure thermal agitation (Fig. 5a-c). Interestingly, supplementing extracts with an energy mix to boost activity enhanced their viscosity, while depleting ATP reduced it (Fig. 1i-j). We interpret these effects from the requirement of energy to sustain F-actin filaments as well as ER anatomy and turnover by for instance promoting membrane fusion and fission^28^. However, in adherent animal cells, depletion of ATP had been rather associated with an increase in viscosity, which was attributed to an ATP-dependent fluidification of the cell interior caused by cytoskeletal molecular motors^45^. These considerations suggest that cellular activity could have opposite influences on cytoplasm physical properties by promoting both the integrity of structural components like endomembranes and cytoskeletal filaments, while enhancing active agitation and consequent fluidification by molecular motors. Future work documenting composite endomembrane-cytoskeletal networks may help discern how the material properties of this class of active matter are regulated.

Eukaryotic cells are filled with many types of vesicular and endomembrane compartments, that come in a range of size, topology and interactions. Yolk granules are specific to early embryos, but pertain to the general class of lipid droplets, that are found in all eukaryotic cells, reaching high packing fractions in specialized cell types like adipocytes, or in the context of several disease states^46^. The ER is also present with similar topology and occupancy in all eukaryotes. As such, our work may have uncovered a universal strategy for eukaryotic cells to control the fluidity of their cytoplasm, by modulating the density, interaction and organization of its endomembranes.

## Lead contact

Further information and requests for resources and reagents should be directed to and will be fulfilled by the lead contact, Nicolas Minc.

## Materials availability

This study did not generate new unique reagents.

## Acknowledgements

We thank all members of the Minc team for discussion and technical help. M.I.A. was supported by a MCSA-PF grant from the EU (“CYTOMECH”-101148446). We acknowledge the ImagoSeine core facility of the Institut Jacques Monod, member of the France BioImaging Infrastructure (https://ror.org/01y7vt929) supported by the French National Research Agency (ANR-24-INBS-0005 FBI BIOGEN) and GIS-IBiSA. This work was supported by the Centre National de la Recherche Scientifique, the Université de Paris, and grants from the Agence Nationale pour la Recherche (ANR, “BulkDev”) to N.M and E.W. and from La Ligue Contre le Cancer (EL2021.LNCC/NiM) and the Fondation Bettencourt Schueller (“Impulscience”) to N.M.

## Author contributions

Conceptualization, N.M., E.W. L.L. P. and M.I.A.; Methodology, M.I.A., A.K., C.M-D, M.M., J.S., S.D., C.D., L-L. P., E.W. and N.M. Writing – Original Draft, N.M. and M.I.A. Draft Editing. M.I.A., A.K., C.M-D, M.M., J.S., S.D., C.D., L-L. P., E.W. and N.M.

## Material and Methods

### Sea urchin gametes

Purple sea urchins (*Paracentrotus lividus*) were obtained from the Roscoff Marine station (France) and maintained at 16 °C in aquariums of artificial sea water (ASW, Reef Crystals, Instant Ocean). Gametes were collected by intracoelomic injection of 0.5 M KCl. Eggs were rinsed twice with ASW, kept at 16°C, and used on the day of collection. Sperm was stored at 4°C and used to verify eggs quality with fertilization tests.

### Sea urchin egg cytoplasm crude extracts

Crude extract (CE) preparation was based on previously established protocols ^34^. Upon collection, unfertilized eggs were gathered together in ASW in a borosilicate beaker and passed several times through a 70 µm Nitex mesh (Genesee Scientific) to remove the jelly coat. The egg suspension was transferred to falcon tubes and centrifuged (Eppendorf Centrifuge 5810 R) for 2 minutes at 200×g to pellet the eggs. The supernatant was removed and eggs were washed 3 times in 10 volumes of lysis buffer (LB) (see below) through repeated resuspension and centrifugation of 2 min at 200×g at 4°C. The exact same amount of supernatant as that of LB was removed to ensure that CE remained undiluted. The pellet was then homogenized in a 15 mL or 7 mL Dounce homogenizer (KIMBLE, DWK Life Sciences), depending on egg volume collected, on ice. This final solution served as undiluted CE (Extended Fig. 1). Varying CE concentrations were obtained by diluting the CE solution in LB, or by evaporating water in a speed vacuum system (Thermo Savant DNA120 Speedvac Concentrator) to overconcentrate extracts.

### Cytosol isolation

Starting from the CE solution, a cytosolic fraction was obtained by following other steps. First, a 5 mL CE solution was centrifuged for 10min at 10,000×g at 4°C (Eppendorf Centrifuge 5425 R) to remove the pellet and the uppermost lipid layer. The middle fraction was then centrifuged at 150,000×g for 3 h at 4°C in a Beckman Optima L-80 XP ultracentrifuge with a SW 41Ti rotor, and the supernatant was collected a membrane-free cytosol extract fraction (Extended Fig. 5) ^34^.

### Yolk granules purification

Sea urchin eggs were collected as described above. Upon jelly coat removal, unfertilized eggs were washed in calcium-magnesium-free artificial sea water (CMF-ASW), and pelleted as low speed^47^. The supernatant was then removed and the pellet was homogenized in 5 volumes of KCl solution (see below) by hand-passing 5 times in the Dounce homogenizer on ice. The yolk granule solution was obtained by repeating twice a two-step procedure. First, the homogenate was centrifuged (Eppendorf Centrifuge 5810 R) for 4 minutes at 400×g at 4°C, and second the pellet was discarded and the supernatant collected and centrifuged for 10 minutes at 2400×g at 4°C. Next, the supernatant was removed and the pellet resuspended in 5 volumes of the KCl solution, and the two steps above were repeated. The final obtained pellet is the yolk granule preparation ^47^. Serial dilution of Yolk suspension, was then achieved by resuspending and diluting the above pellet in membrane-free cytosol.

### ER purification

Starting from the CE solution, the ER purified solution was obtained through small modifications of the protocol from Leiro et al^48^ (Extended Fig. 6). Sequential centrifugation steps were performed to progressively clarify the solution. All steps were done at 4°C. First, 5mL of the CE solution was centrifuged at 600×g for 5 minutes to remove debris. The pellet was discarded, and the supernatant was centrifuged at 600×g to further cleanse the sample. Then, a series of centrifugation steps was performed in an Eppendorf Centrifuge 5425 R and the pellets were systematically discarded and the supernatant kept:1,200×g for 4 minutes, 7,000×g for 15 minutes, and 20,000×g for 35 minutes. The final supernatant was then centrifuged at 100,000×g for 65 minutes to collect ER membranes, using a Beckman Optima L-80 XP ultracentrifuge with a SW 41Ti rotor. This ER pellet was resuspended in 150µL cytosol but still contained a large fraction of yolk granules, so it was characterized and used as ER/yolk fraction. The pure ER fraction was obtained through an additional centrifugation step at 20,000×g for 35 minutes (Eppendorf Centrifuge 5810 R) to segregate yolk granules in the pellet and the obtained supernatant was used as purified ER solution in cytosol.

### Buffers and solutions for extract fraction preparations

The Lysis Buffer (LB) consists of 10 mM HEPES pH 8.0, 250 mM NaCl, 25 mM EGTA, 5 mM MgCl2, 110 mM Glycine, 250 mM Glycerol, and 1 mM DTT and 1 mM PMSF were added just before use it. Calcium- and magnesium-free artificial see water (CMF-ASW) were prepared by mixing 7.1g of NaCl, 0.2g of KCl, and 2.1g of Na_2_SO_4_ in 250mL of DI Water. The KCl solution used for yolk purification consisted of 0.55M KCl and 1mM EDTA at pH 7.0. Fixative Buffer consists of 0.1M (pH7.2) sodium cacodylate trihydrate, 2%v/v of sucrose, 2.5%v/v of glutaraldehyde (25%), and 2%v/v of paraformaldehyde (16%). The energy mix was obtained by combining 2mM ATP, 1mM of GTP, 2mM MgCl2, and 15mM creatine phosphate.

### Chemical inhibitors

F-actin inhibition was achieved by adding 0.5µL of cytochalasin D (Sigma-Aldrich) to 50µL of CE solution to reach a final concentration of 20 μM starting from a 1mg/mL stock in DMSO. 15 minutes later, a representative sample of 5 µL was taken and stained with Alexa-Phalloidin to visualize F-actin (see below). For rheology experiment, 3 µL of cytochalasin D were added to 300µL of CE solution. ATP depletion was performed by supplementing CE with 2 mM NaN_3_ (Sigma-Aldrich) and 10 mM 2-deoxy-d-glucose (Sigma-Aldrich). After 1 hour of incubation the sample was analyzed under the microscope. In the case of rheological assays, the solution was taken for preparing the droplet encapsulation after 30 minutes of incubation, and analyzed at least after a total time of 1 hour.

### Fluorescent labeling of intracellular components

To label mitochondria, 5mL of eggs were incubated for 20–30 minutes with 2.5 µL of MitoTracker (Thermo Fisher), and a CE suspension was prepared from these pre-labelled eggs. Yolk granules were labeled directly in 5 µL CE (likewise, in yolk granule an ER solutions) by adding 0.5µL from a 2 mg/mL stock of Nile Blue or Nile Red diluted in DMSO (Sigma). ER membranes staining was achieved using a saturated solution of DilC18(3) in oil prepared by mixing several crystals of DilC18(3) (Thermo Fisher) in 100 μL of soybean oil. A 0.5µL of DilC18(3) solution was added to 5µL of CE (likewise, in yolk granule and ER solutions). Alexa Fluor 488 phalloidin (Invitrogen) at 1 U per 100 μL in LB was used to stain F-actin by adding 0.5-1µL to 5µL of CE (likewise, in yolk granule an ER solutions). 0.5µL of purified Tau-mBD-StayGold (1mg/mL, stock concentration) was added to 5µL of CE to label Microtubules.

### Imaging

Time-lapses of beads moving under magnetic force were recorded on an inverted microscope equipped with a micromanipulator for magnetic tweezers, at a stabilized room temperature (18–20°C). The set-up consisted of a Leica DMI6000 B microscope, operated with Micro-Manager (Open Imaging), equipped with HCX FL PLAN 10x/0.25 Ph1 objective (Leica), and a HC Plan APO 20x/0.70 objective (Leica), and an ORCA-Flash4.0LT (Hamamatsu) camera. Imaging was done in DIC/fluorescence at rates varying from 4 frames to 1 frame per second.

To visualize CE and other extract fractions, live imaging was performed on three different microscopes. A Nikon Ti-Eclipse microscope equipped with an ORCA-Flash4.0LT (Hamamatsu) camera, using a 100X oil-immersion objective (Plan Apo VC, Nikon), and operated with Micro-Manager (Open Imaging). A Nikon Ti-Eclipse microscope, equipped with a CSU-X1 (Yokogawa) confocal spinning head, and a Prime BSI (Teledyne Photometrics) camera, using a100x oil-immersion objective (CFI Plan Apo 100X Oil, Nikon), and operated with Metamorph software (Molecular devices). A Nikon Ti2-Eclipse microscope, equipped with a CSU-W1 (Yokogawa) confocal spinning head, and an Kinetix 22 (Teledyne Photometrics), using a 100X oil-immersion objective (CFI Plan Apochromat Lambda D 100X Oil, Nikon), and operated with VisiView software.

### Serial Block Face Scanning Electron Microscopy

Details of endomembranes organization were captured using serial block face scanning electron microscopy (SBF-SEM). Unfertilized eggs were fixed in 0.2M sodium cacodylate, 0.25M sucrose, 2% paraformaldehyde, 2% glutaraldehyde (EM Fixative Buffer) for 1h at room temperature. Fixed sample were then progressively transferred in 0.2M sodium cacodylate, 0.35M NaCl. Samples were further processed for SBF/SEM following the NCMIR protocol (https://ncmir.ucsd.edu/sbem-protocol) as previously described^19^. Processed samples were mounted on aluminum pins, trimmed and inserted into a TeneoVS SEM (Thermo Fisher Scientific). 3D imaging using back-scattered electrons was performed at 2.7kV with a beam current of 400pA, a dwell time of 1μs per pixel under 40Pa vacuum. The thickness of serial sections was 100nm.

### Transmission electron microscopy

CE and purified Yolk granules solutions were fixed by incubation in the EM Fixative Buffer (1:1) for 15 min, pelleted to remove the fixative, washed with CE-buffer and centrifuged again. The pellets were embedded in 1% agarose by mixing (CE or Yolk granules) with 2% agarose at a 1:1 ratio. The samples were cut into small 1 mm3 pieces, and then transferred into PBS (pH 7.4). Post-fixation was performed in 1% OsO4 reduced with 1.5% potassium ferrocyanide in PBS for 1h at room temperature. After washing, dehydration was carried out using graded concentrations of ethanol in water for 10 min each. Resin infiltration was performed for 2h in a 30% Agar low viscosity resin (Agar Scientific Ltd), then overnight in 50% resin, followed by three 2h-incubations in pure resin, prior to inclusion in BEEM capsules and 18 hours polymerization at 60 °C. 70 nm ultrathin sections were obtained using an EM UC6 ultramicrotome (Leica), and post-stained in 4% aqueous uranyl acetate followed by lead citrate (Reynold’s solution, deltamicroscopies, France). Sections were examined with a 120 kV TEM (Tecnai12, Thermo Fischer Scientific) equipped with a 4K CCD camera (Oneview,Gatan).

### Segmentation of endomembranes

Optical microscopy images were processed using Ilastik a machine-learning-based interactive tool (https://www.ilastik.org/) . The Pixel classifier was trained iteratively with the corresponding label (ER, yolk, or mitochondria) until a satisfying accuracy was reached for endomembrane compartments. Pixel classification maps were then extracted from the full-scale dataset. For SBF-SEM images, Yolk granules, mitochondria and endoplasmic reticulum were segmented from the resulting image stacks with a supervised learning approach using the Ilastik software (1.4.0-gpu)^49^. Pixel classifications were obtained through an independent autocontext workflow for each compartment. 3D renders were generated from the pixel probability maps in Blender using the *microscopynodes* add-on^50^.

### Quantification of particle distribution heterogeneity

The spatial heterogeneity of the particle distribution was quantified from the coordinates of particle centers extracted from microscopy images. For each image, the particle coordinates were extracted from segmented images obtained with a supervised learning approach using the Ilastik software (1.4.0-gpu)^49^. Image fields were divided into a regular square grid covering the entire field of view. The grid cell size was fixed to 10 µm × 10 µm, providing a compromise between spatial resolution and statistical sampling. To minimize boundary effects, the outermost row and column of grid cells were excluded from the analysis. For each remaining grid cell, the local particle density was calculated as the number of particle centers contained within the cell divided by the cell area. The spatial heterogeneity of the suspension was then quantified using the coefficient of variation of the local particle density (C_V_), defined as the standard deviation of the local particle density divided by its mean value: C_V_ = σρ /⟨ρ⟩, where ⟨ρ⟩ is the mean local particle density and σρ is its standard deviation over all analysed grid cells. A homogeneous particle distribution yields low values of C_V_, whereas the formation of dense aggregates separated by particle-depleted regions leads to an increase in C_V_, reflecting the increasing spatial heterogeneity of the suspension. The analysis was performed in MATLAB (MathWorks) using the same analysis parameters for all images. To ensure that the measured trends were not dependent on the chosen spatial discretization, the analysis was repeated using different grid sizes (5–20 µm), yielding consistent qualitative results.

### Droplet fabrication

Water-in-oil cell-like compartments droplets were fabricated based on a previously established capillary-trap setup ^26^. The solution of interest (CE, Yolk, ER, ER+Yolk) supplemented with magnetic beads was placed in a 1mL syringe (Terumo) connected to a PVC microfluidic tube through a fine needle (Hamilton Kel-F Hub Needles). The free end of the tube was connected to a fused silica capillary (Polymicro Technologies LLC) tip of 100.6 µm internal diameter that is periodically moved through an air-oil interface thanks to a 3D printer motorized arm. The oil solution is contained in a 50 mm Glass Bottom dishes (MaTek Corporation). As the tube crosses the air-oil interface, the aqueous droplet at its tip detaches. The oil phase is composed of a non-ionic surfactant, Span 80 (Sigma Aldrich), dispersed in an equal mix (1:1) of silicon oil AR20 (Sigma Aldrich) and pure hexadecane (Sigma Aldrich) at a mass concentration of 2% (w/w). Droplets containing the solution of interest and one agarose magnetic bead were used for active rheology assays. Once fabricated, droplets were placed inside hollow square capillaries (ID 500 µm, Vitrocom Inc.) for magnetic force application ^27^.

### Magnetic force calibration

Magnetic forces were calibrated *in vitro* following procedures described previously^19,51,52^. The magnetic force field created by the magnet tip was first characterized by pulling super-paramagnetic 1 µm Dynabeads (MyOne Streptavidin C1; Thermofisher) in a viscous test fluid (80% glycerol, viscosity 8.0×10−2 Pa sec at 22°C) along the principal axis of the magnet tip. Small motion of the fluid was subtracted by tracking 0.5-μm nonmagnetic fluorescent tracers (Molecular probes; Invitrogen) in the same suspension. The speed of the magnetic beads (*v*) was computed as a function of the distance to the magnet, to obtain and trace the decay function of the magnetic force, which was fitted using a double exponential function^51,52^.

To compute the dependence of the force on the size of the bead, agarose beads (PureCube Glutathione MagBeads, Cube Biotech) of diverse sizes were pulled in the same fluid as above. The speed (*v_a_*) was measured and translated into a force using Stokes’ law *F=6πηRv_a_*, where *η* was the viscosity of the test fluid and *R* the bead radius. The force–size relationship at a fixed distance from the magnet was well represented and fitted by a cubic function. These speed–distance and force–size relationships were combined to compute the magnetic forces applied to the agarose beads as a function of time, from the size of the beads and their distance to the magnet tip.

### Magnetic force application

Magnetic tweezers were implemented as described previously ^16,51^. The magnet probe used for force application was built from three rod-shaped strong neodymium magnets (diameter 4 mm; height 10 mm; S-04-10-AN; Supermagnet) prolonged by a sharpened steel piece with a tip radius of ∼50 μm to create a magnetic gradient. The surface of the steel tip was electro-coated with gold to prevent oxidation. The probe was controlled with a micromanipulator (Injectman 4, Eppendorf), mounted on an inverted microscope. Agarose beads (PureCube Glutathione MagBeads, Cube Biotech) of diverse diameters were used to apply magnetic forces in cytoplasm extracts.

### Tracking of bead position

Magnet tip position and magnetic bead displacement was recorded in DIC for droplets placed directly in the oil container or in capillaries. Agarose beads time-lapse movie color was inverted to track the position of the bead using the TrackMate plugin in Fiji^41^ . The trajectories of the beads were rotated to align their displacement vector parallel to the horizontal x-axis. For droplets analyzed directly in the oil container, small displacement of the droplets could occur from the insertion of the magnet tip in the solution, this shift in the position of the droplets was corrected by translate transformation in Fiji.

### Viscoelastic parameter calculation

Magnetic bead displacements were fitted with a Jeffreys’ model using a custom written code in Matlab (Mathwork) to compute viscoelastic parameters. For the rising phase, the position was fitted using:

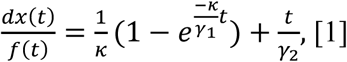

where *dx* is the displacement along the x-axis and *f* is the magnetic force. This rescaling of the displacement by force allows to compensate for variations in force amplitude during each pull, and implicates that we assume that viscoelastic responses are mostly linear. These fits allowed to compute the restoring stiffness, *κ*, and the viscoelastic drags *γ*_1_ and *γ*_2_ of oil droplets and beads, and thus the viscoelastic timescales as *τ*_1,2_ = *γ*_1,2_/*κ* . Stiffness and drags were converted into bulk elastic modulus and shear viscosities using particle size and generalized Stokes law. Trajectories of beads during the relaxation phase were influenced by the active cytoplasm extract fluctuations so that beads often exhibited random motions after moving a few steps backward. As such we restricted the analysis of viscoelastic relaxation to a phase before Brownian motion started to dominate characterized by an angle between two successive steps exceeding 90°. Relaxation phases were fitted using:

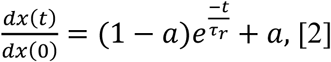

where *t = 0* corresponds to the time of the end of force application, and *τ*_r_ is the timescale of decay. Fits of the obtained data for every single experiment were done using the optimization method in Matlab in order to minimize sensitivity to initial fitting values.

### Mean Square Displacement (MSD)

To track yolk in CE and in the ER/yolk suspension, yolk granules were stained with NB as described above. For the pure yolk suspension, yolk purified solution was diluted in cytosol to reach a volume fraction similar to that *in vivo* an in CE, and was also stained with NB. Imaging was done under a Nikon Ti2-Eclipse microscope, operated with VisiView software and equipped with a CSU-W1 (Yokogawa) confocal spinning head, and an Kinetix 22 (Teledyne Photometrics), using a 100X oil-immersion objective (CFI Plan Apochromat Lambda D 100X Oil, Nikon) at rates of 300ms. The TrackMate plugin in Fiji was used to track the position of yolk granules. After that, the obtained trajectories were analysed with a Matlab home-made code to compute MSD curves. The curves were fitted to compute the diffusion coefficient, *D*, and the power law, *α*, using the equation *MSD* = 4*Dτ*^a^. D was computed by fitting the averaged MSD in a linear scale using a fraction of points covering short time-scale on which the MSD was linear with lag time, while α was computed by a linear fit of the whole of averaged MSD in log-log scale (Extended Fig. 8b-c).

### Flow analysis

Time-lapses of creep assays were used to analyze flows in cell-like compartments with the particle image velocimetry PIVlab tool in Matlab. The droplet exterior was masked to be excluded from the analysis. Contrast limited adaptive histogram equalization and two-dimensional Wiener filter were applied on the images in the pre-processing steps for noise reduction (with accordingly windows of 20 and 3 pixels widths). Image sequences were investigated in the Fourier space by four interrogation windows with 64, 32, 16, and 8 pixels widths and 50% overlapped area. The spline method was used for the window deformation and subpixel resolution obtained by two-dimensional Gaussian fits. After smoothing, the output vector fields were used for further analysis and for plotting flow maps and streamlines in Matlab.

## Statistical analysis

All experiments presented in this MS were repeated at least twice and quantified using a number of cell-like compartments or events reported in each figure panel legend.

## Simulation

We implemented and performed agent-based simulations to assess the role of endomembrane crowding in determining cytoplasm viscoelasticity. The simulations are based on a highly coarse-grained description of the cytoplasm and were implemented in *Cytosim*, an open-source package based on overdamped Langevin dynamics^43^ . The simulations explicitly consist of two main agents: 1-Suspensions of spherical beads representing organelles such as mitochondria and yolk granules. Each particle has a diameter of 1µm, corresponding to the typical size of yolk particles in sea urchin embryos. A steric repulsion between particles was implemented to prevent overlap and to generate excluded volume interactions. 2-Assemblies of elongated filaments interconnected by crosslinkers, mimicking percolated tubular membranous endoplasmic reticulum (ER). The filaments are non-extensible and discretized into N segments. The bending energy per unit length of the filament is given by ½ κ C^2^, where κ and C are the bending rigidity and curvature of the filament, respectively. Filaments also experience steric interactions both with one another and with the spherical particles. To generate a connected ER meshwork, four crosslinkers were assigned per filament and modeled as elastic springs of stiffness k_cr_. Crosslinkers bind along filaments within a binding range, r_b_, at a constant binding rate ω_on_. Bound crosslinkers can detach from the filament with an off-rate ω_off_, that depends on both time and the resistive load: ω_off_= ω_d_ exp(lfl/fd), where ω_d_ is a constant detachment rate and f_d_ is the characteristic unbinding force. In general, the unbinding rate was set to relatively high values, making crosslinkers mostly stable over the time course of simulations. All components were confined within a spherical domain mimicking the cell membrane or cell-like compartments. The background viscosity was implemented as to regulate the mobility of moving particles, and was set equal to that of measured membrane-free cytosol viscosity. Finally, a probe particle of radius 8 µm was embedded in these synthetic cytoplasms. A constant force was applied to the probe to induce a displacement and was released after 3.5 s, reproducing active magnetic-tweezers microrheology experiments. This configuration corresponds to a 100% cytoplasm concentration and dilution can be simulated by reducing the density of the agents. A complete list of simulation parameters is provided in Table S1.

**Table S1.** Simulation parameters and conditions.

| Symbol | Parameter | Value | Notes |
| --- | --- | --- | --- |
| $k_B T$ | Thermal energy | 4.2 pN nm | Room temperature |
| $\eta_0$ | Background viscosity | 0.02 pN.s/$\mu\text{m}^2$ | Cytosol viscosity |
| $\Delta t$ | Time step | 0.005 s | |
| $R_{\text{cell}}$ | Cell radius | 30 $\mu\text{m}$ | |
| $k_s$ | Strength of steric repulsion | 200 pN/$\mu\text{m}$ | Varied in Extended figure 4b |
| **ER Filaments** |  |  |  |
| $L$ | Filament length | 6 $\mu\text{m}$ | |
| $d_0$ | Filament diameter | 0.2 $\mu\text{m}$ | $2 \times$ ER tubes |
| $EI$ | Filament stiffness | 0.04 pN/$\mu\text{m}^2$ | |
| $N_{\text{ER}}$ | Number of filaments | 43200 | 7.2 % volume fraction |
| $ds$ | Filament segmentation | 0.2 $\mu\text{m}$ | |
| **Yolk Beads** |  |  |  |
| $r$ | Yolk radius | 0.5 $\mu\text{m}$ | |
| $N_y$ | Number of yolks | 64800 | 30 % volume fraction |
| **Probe** |  |  |  |
| $R$ | Probe radius | 8 $\mu\text{m}$ | |
| $F$ | Force inserted to the probe | 500 pN | |
| **Crosslinkers** |  |  |  |
| $N_{\text{Cr}}$ | Number of crosslinkers per filament | 4 | Varied in Extended figure 4a |
| $\omega_{\text{on}}$ | Binding rate | 30 $\text{s}^{-1}$ | |
| $r_b$ | Binding range | 0.5 $\mu\text{m}$ | |
| $\omega_{\text{d}}$ | Unbinding rate | 1 $\text{s}^{-1}$ | |
| $f_{\text{d}}$ | Unbinding force | 50 pN | |
| $k_{\text{cr}}$ | Link stiffness | 100 pN/$\mu\text{m}$ | |

## Extended Figures

**Extended Figure 1.**
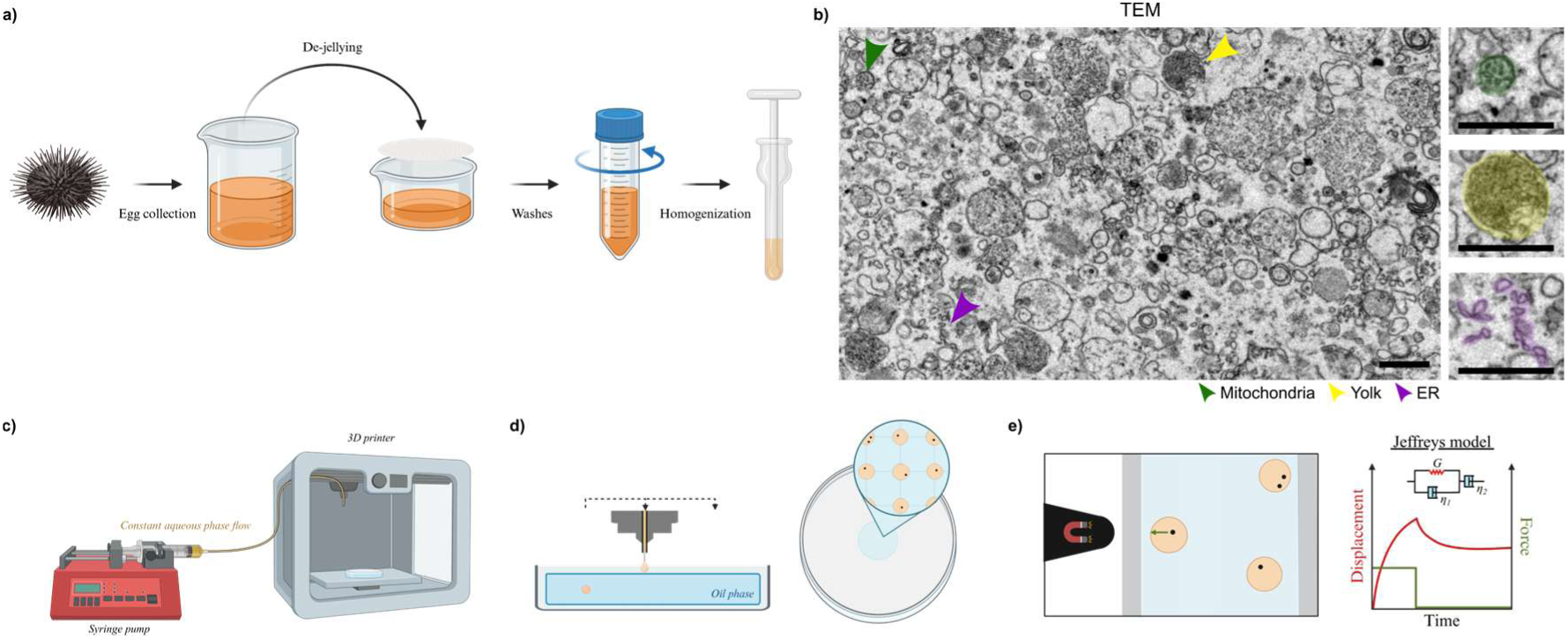
Cytoplasm extracts preparation, cell-like compartments and droplet fabrication for active microrheological measurements. **a**, Eggs from *Paracentrotus Lividus* animals are collected in ASW after intracoelomic injection of animals in gonads, and filtered through a mesh to remove the jelly coat. Eggs are then centrifuged, dried and washed and the final pellet of eggs is homogenized using a hand-held Dounce homogenizer, to generate nearly-undiluted crude extracts (CE). **b,** Representative transmission electron microscopy image of CE. The arrows of different colors point to particular cytoplasm endomembrane components in the CE, and the insets highlight supervised-learning based registration of different endomembranes. Scale bar: 1µm. **c,** Syringe pump and 3D printer setup generating a constant flow of the aqueous phase containing magnetic beads. **d,** The needle tip follows a grid-like motion, periodically dipping into an oil layer to generate mono-disperse and regularly distributed aqueous droplets. Droplets containing magnetic beads are deposited in a spatially organized array within the oil. **e,** A magnet tip actuated by a micro-manipulator serves to apply a calibrated magnetic force (green arrow) on the bead inside the droplet. The droplet is held in a capillary to prevent droplet motion. Bead displacement in response to applied force is modelled using Jeffreys’ viscoelastic model, with the force step shown in green and resulting displacement of the bead in red.

**Extended Figures 2.**
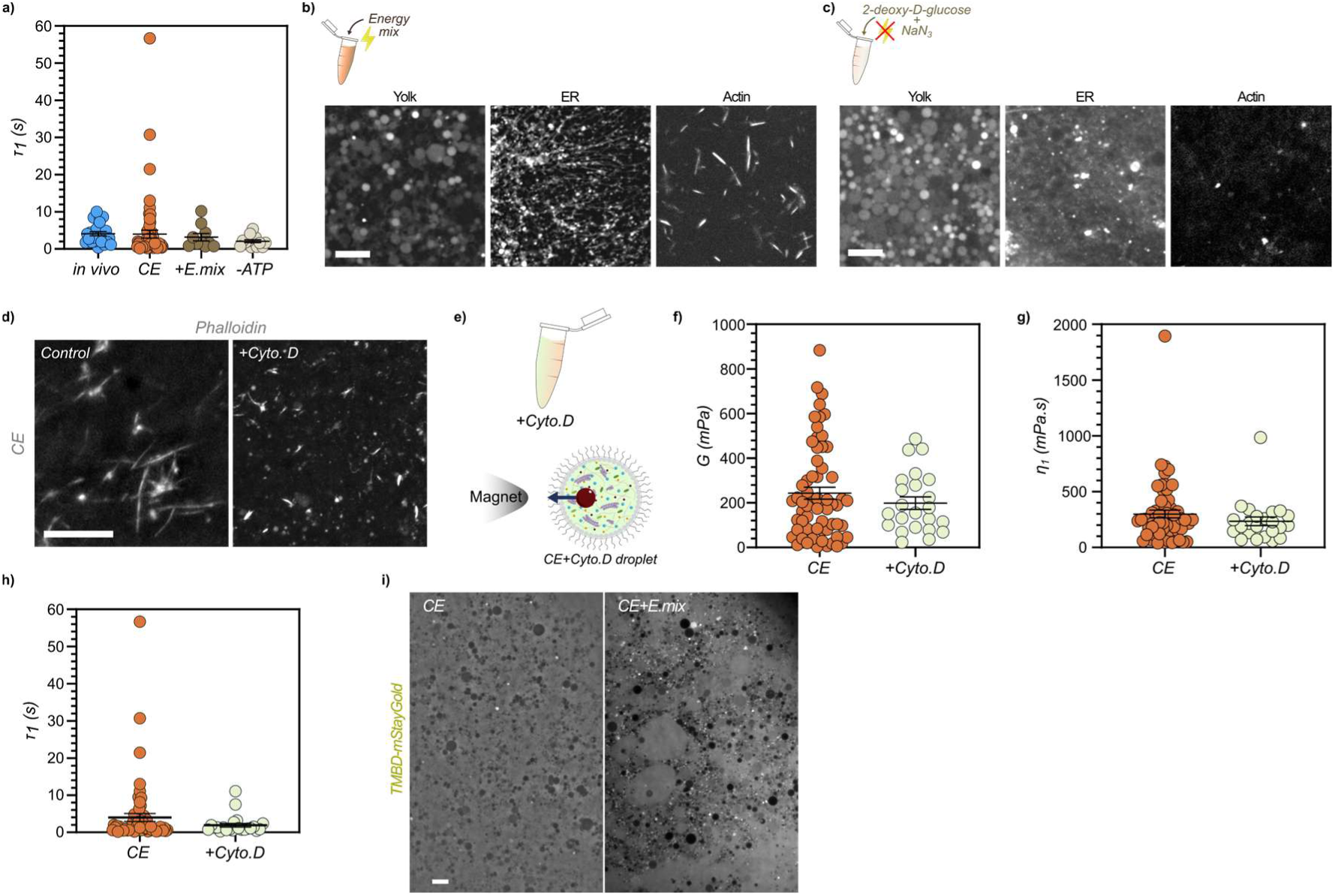
Contribution of the cytoskeleton and chemical energy to CE viscoelasticity. **a**, Viscoelastic timescale obtained from fitting creep curves with Jeffreys’ model for in vivo cytoplasm^16^(n=23), CE (n=62), CE supplemented with an energy mix solution (n=10), and ATP-depleted CE solution (n=14), corresponding to values obtained in Fig.1 i and j. **b,** Spinning disk confocal images of CE supplemented with an energy mix and stained for yolk (Nile Blue), ER (DilC18), and F-actin (Phalloidin). Scale bar: 5µm. **c,** Spinning disk confocal images of CE supplemented with 2mM sodium azide and 10mM deoxyglucose to deplete ATP and stained for Yolk (Nile Red), ER (DilC18), and F-actin (Phalloidin). Scale bar: 5µm. **d,** Spinning disk confocal images of CE and CE treated with cytochalasin D and stained for F-actin. Scale bar: 10 µm. **e,** Scheme of Cytochalasin D treated CE encapsulated with magnetic beads for active rheology. **f,g,** Elastic and viscous moduli of control CE and CE treated with cytochalasin D (n=62 and n=23, respectively). **h,** Viscoelastic timescale for CE and CE treated with cytochalasin D D (n=62 and n=23, respectively). **i,** CE solution stained with TBMD-mStayGold to label Microtubules showing the absence of microtubules in control conditions and in CE solution with an energy mix. Scale bar: 5µm.

**Extended Figures 3.**
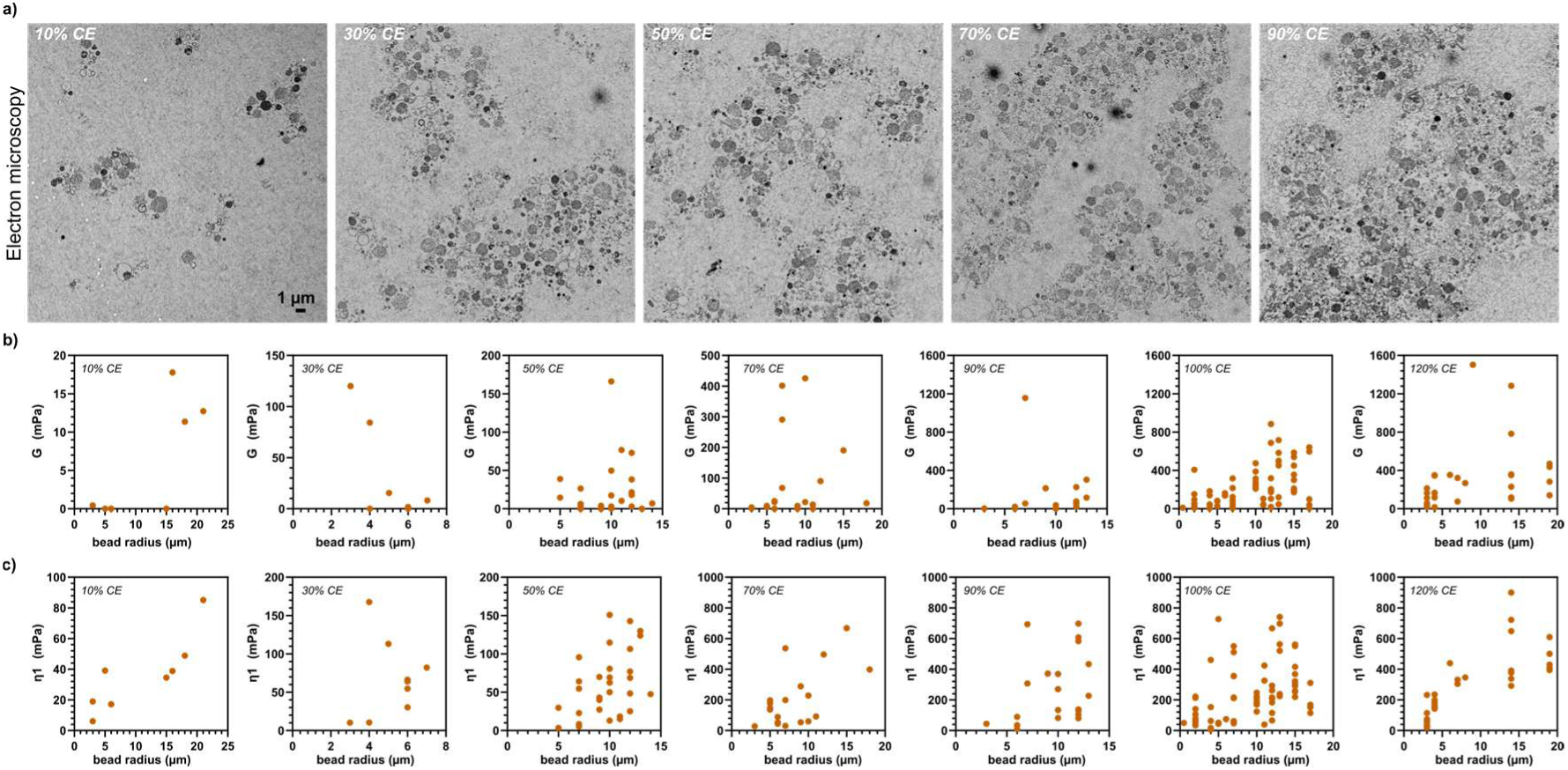
Study of the crude extracts at different dilutions. **a**, Transmission Electron microscopy images of CE diluted at different concentrations. Scale bar: 1µm. **b,c,** Viscoelastic constants as a function of bead size for different CE concentrations.

**Extended Figures 4.**
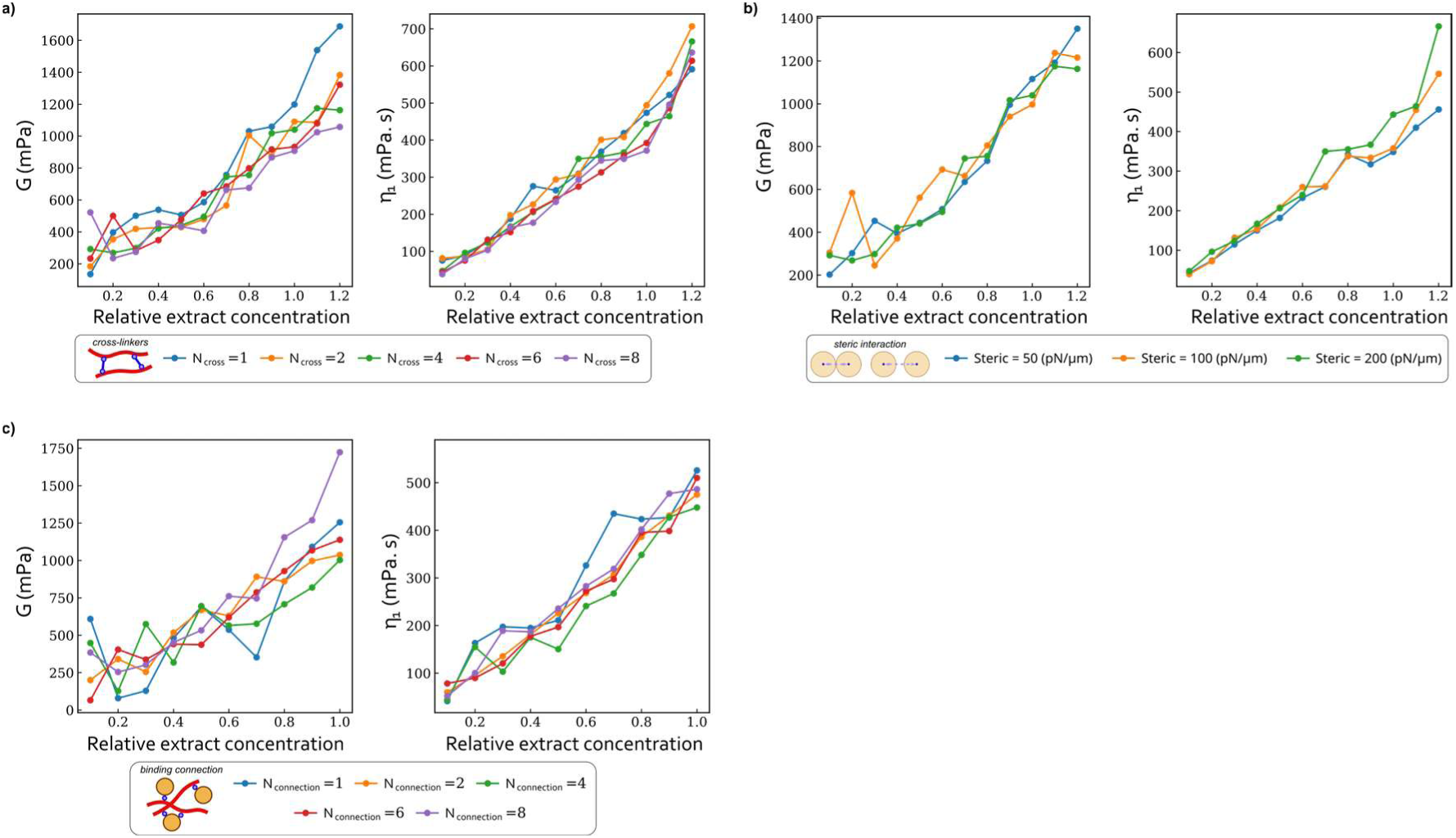
Impact of simulation parameters on viscoelastic constants in the model. **a-d**, Simulation of the elastic and viscous constants at different extract concentration to explore the effect of parameters for (a) number of crosslinkers per filament, (b) steric interactions among spherical particles, (c) connections between yolk particles and ER filaments.

**Extended Figure 5.**
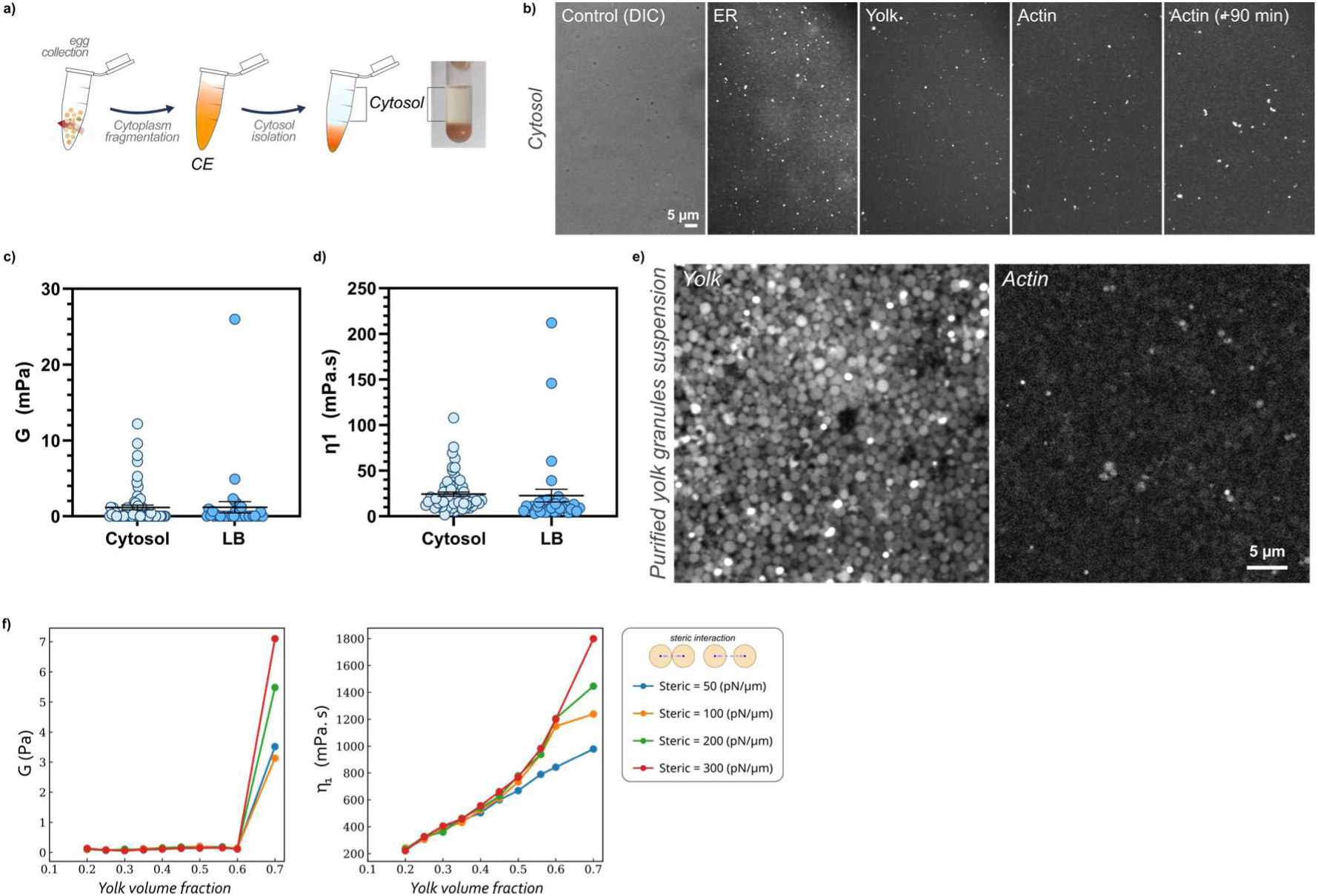
Decomposing the cytoplasm. **a**, After egg collection, crude extracts (CE) were obtained and layered by ultracentrifugation to isolate a membrane-free cytosolic fraction. **b,** Cytosol fraction labelled for ER, Yolk and F-actin validating the absence of endomembrane and cytoskeleton networks in cytosolic fractions. **c,d,** Elastic (c) and viscous (d) moduli measured for the cytosol and LB mediums with magnetic tweezers based rheology in cell-like compartments D (n=64 and n=35, respectively). **e,** Purified yolk granule suspension stained with Nile Blue to visualize yolk and with Phalloidin to visualize F-actin, showing the absence of F-actin filaments. **f,** Simulation results for different steric interaction values of the viscoelastic constant predicted for a range of yolk granules volume fractions.

**Extended Figure 6.**
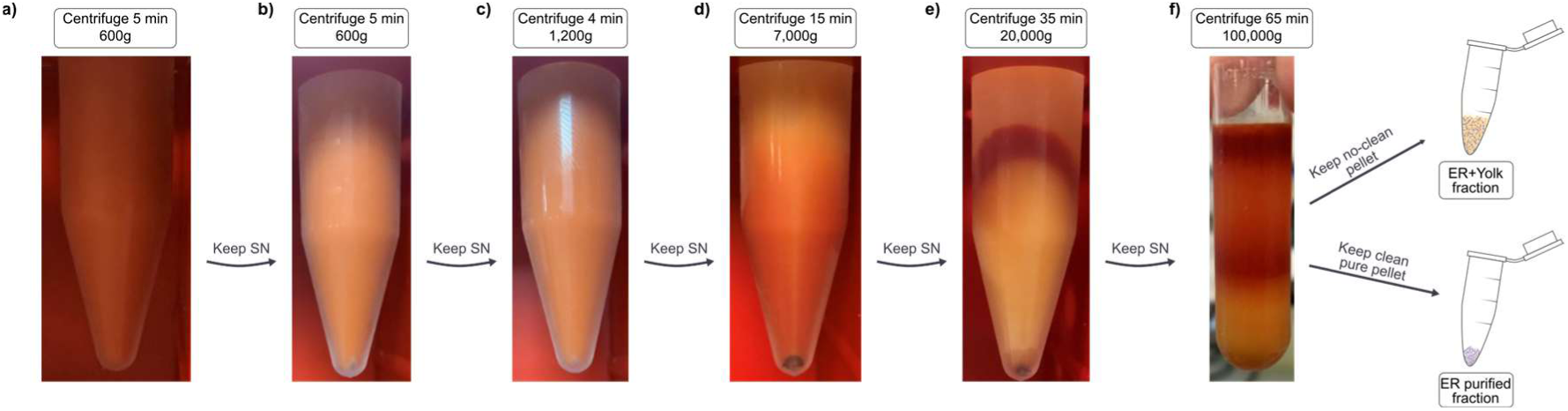
Purification of ER and ER+yolk fractions. **a**, Starting from a 5mL solution, CE were centrifuged for 5 minutes at 600×g at 4°C, the pellet was then discarded and the supernatant (SN) kept. **b,** The SN was centrifuged for 5 minutes at 600×g at 4°C, the pellet was discarded and the supernatant (SN) kept. **c,** The SN was centrifuged for 4 minutes at 1,200×g at 4°C, the pellet was discarded and the supernatant (SN) kept. **d,** The SN was centrifuged for 15 minutes at 7,000×g at 4°C, the pellet was discarded and the supernatant (SN) kept. **e,** The SN was centrifuged for 35 minutes at 20,000×g at 4°C, the pellet was discarded and the supernatant (SN) kept. **f,** The SN was centrifuged for 65 minutes at 100,000×g at 4°C, the two pellets obtained gives a ER+yolk fraction and a pure ER fraction.

**Extended Figure 7.**
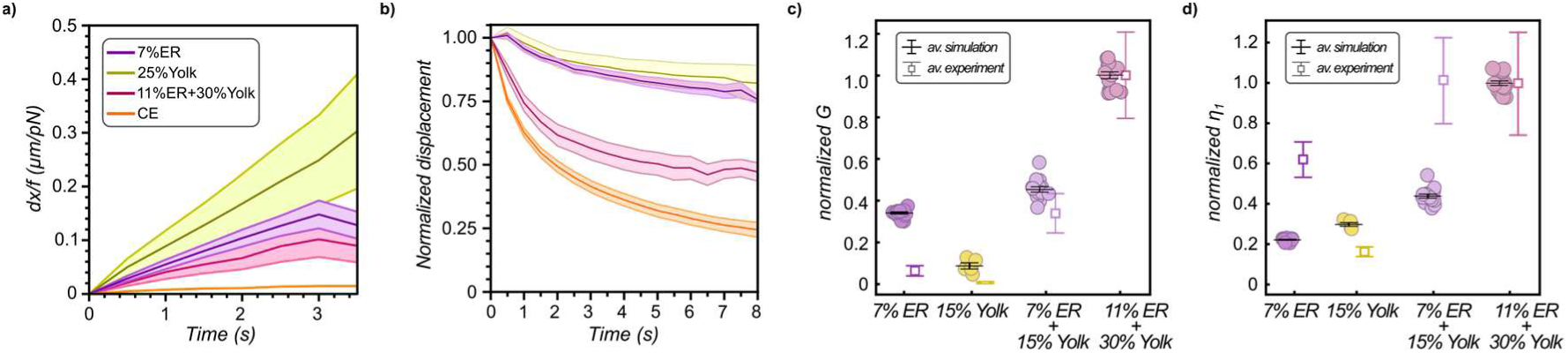
Viscoelastic behavior of composite endomembrane suspensions in experiments and simulations. **a,b**, Creep (a) and relaxation (b) curves for different extract fractions: 7%ER, 25%Yolk, 11%ER+30%yolk, and CE (n=15, n=5, n=18, and n=20, respectively). **c,d,** Normalized elastic (c) and viscous (d) moduli for simulation and experiments for different extract fractions: 7%ER, 25%Yolk, 11%ER+30%yolk, and CE (n=5 independent simulations). Error bars and shades represent +/- s.e.m.

**Extended Figure 8.**
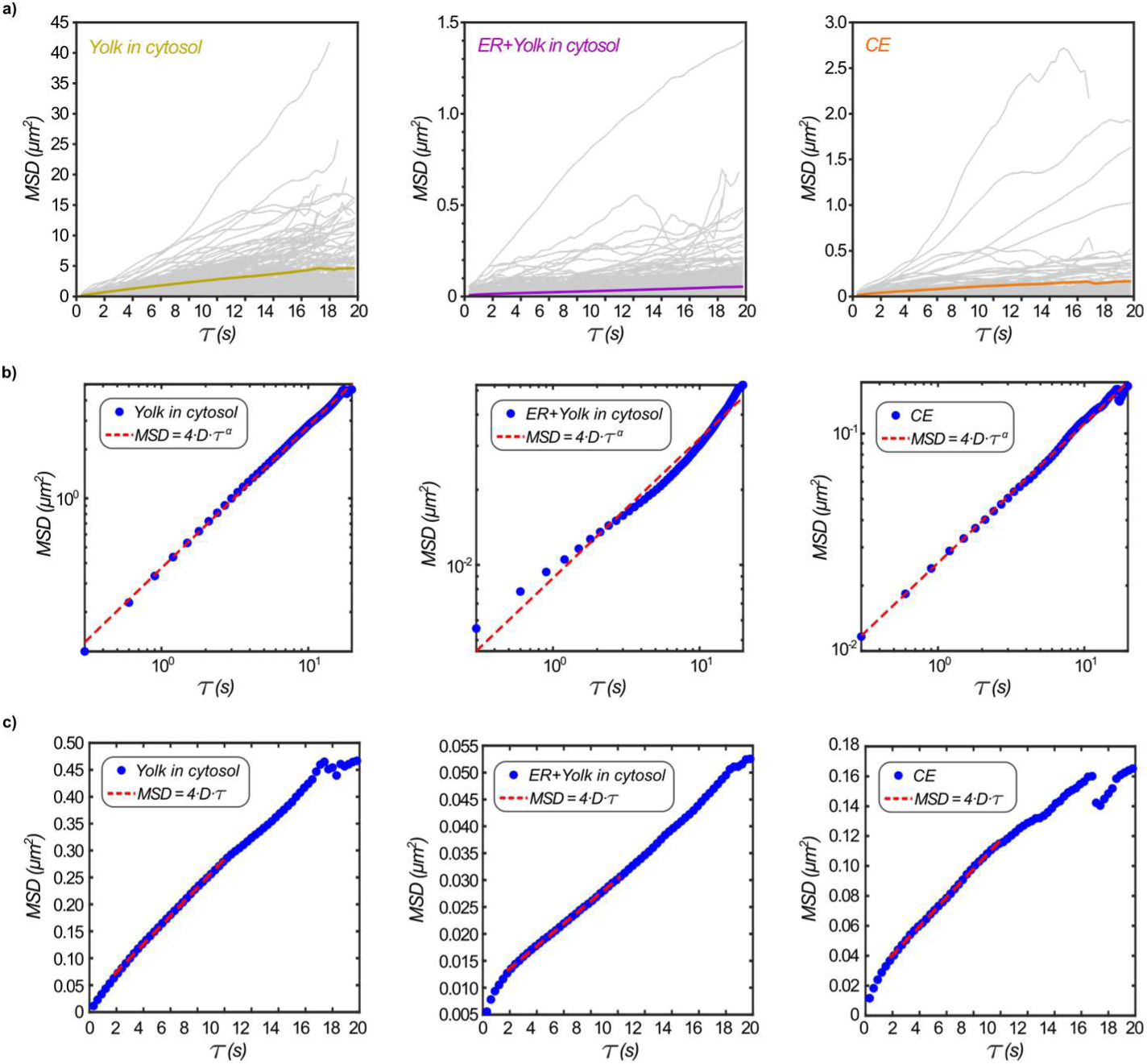
MSD analysis of yolk diffusion in different experimental conditions. **a**, Individual MSD traces for yolk granules tracked in cytosol (left), yolk in a yolk+ER suspension in cytosol (middle), and yolk in CE (right). **b,c,** Average MSD curves in log-log or normal coordinates, used to compute α (b) and D (c) by fitting the specified curves, as indicated by the red dotted line.

### Supplemental Movie Legends

**Movie S1.** 3D reconstitution and rendering of a cytoplasm cube of 2um*2uml*2um obtained upon segmentation and registration of endomembranes from SBF-SEM imaging.

**Movie S2.** Live imaging of crude extracts stained with Dil Dye to label the endoplasmic reticulum.

**Movie S3.** Example of a magnetic particle pulled inside a cell-like compartment filled with crude extracts.

**Movie S4.** Live imaging of the endoplasmic reticulum labeled with Dil Dye during a magnetic pull. Note the deformation of membranes at the bead front.

**Movie S5.** Agent based simulation of creep and relaxation response in a model cytoplasm. (Left) Endoplasmic reticulum, (Middle) Yolk granules, (rigth) Merge.

**Movie S6.** Model cytoplasm simulations at different dilutions, and corresponding creep and relaxation responses below. Note that Y-axis have different ranges.

**Movie S7.** Yolk suspension simulations at different dilutions, and corresponding creep and relaxation responses below. Note that Y-axis have different ranges.

